# A novel tool for monitoring endogenous human MYC transcription and translation by EGFP tag insertion at the 3’ end using CRISPR-Cas9 genome editing

**DOI:** 10.1101/603746

**Authors:** N. Cumbal, MD. Cole

## Abstract

The MYC oncogene is overexpressed in over 70% of human cancers. Since its identification, the study of MYC has led to the discovery of the various ways through which oncogenes contribute to the ability of normal cells to become malignant. However, there are many aspects of MYC biology that remain unknown or controversial in terms of its regulation, targetability and downstream control of its targets. We developed two stable cell lines expressing MYC endogenously tagged with EGFP via CRISPR/Cas9-mediated genome editing. This system allows efficient detection of transcriptional activity of MYC as well the resulting fusion protein while maintaining the gene expression profiles, growth factors-associated MYC induction and growth kinetics of the parental cells. To our knowledge, this is the first report showing endogenous monitoring of MYC expression in colorectal adenocarcinoma through an EGFP tag, thus making it an efficient tool for high-throughput approaches such as genetic and drug screens.

## Introduction

MYC is an oncogene whose expression is associated to the entry of cells into the cell cycle and cell proliferation^1,2^. Due to these fundamental functions and the role MYC plays in most biological phenomena, MYC expression is finely regulated at the transcriptional and translational level. Dysregulation of MYC expression through various mechanisms such as chromosomal translocations, aberrant signaling, amplification and mutations causes normal cells to transform into malignant cells^3,4,5,6^. MYC is documented to play a role in tumor initiation by inhibiting cell differentiation and promoting the reprogramming of differentiated cells^7,8^. Interestingly, the depletion of MYC in established cancer cell lines uniformly reduces cell proliferation and even drives apoptosis, suggesting that MYC is a prime therapeutic target^9,10,11^. A few experimental models for MYC characterization have been developed. Among these, cell lines containing the inducible MYC-ER fusion construct^12^ and mouse models containing N-terminal EGFP-MYC^13^ fusion protein have been used to shed light on topics such as the regulation of MYC and its downstream regulation of target genes. Similarly, many of the known aspects of MYC biology were characterized by using exogenous reporters ^14,15,16^ that don’t take into account the presence of vast genomic regions surrounding the MYC locus that harbor regulatory potential through epigenetic mechanisms^17^. Indeed, advances in our understanding of epigenetic architecture and its effect on transcriptional regulation^18^ indicate the importance of innate chromatin structure in determining the complex interactions of transcription factors, epigenetic factors and cis-acting elements. The development of CRISPR-mediated gene editing technology and its ability to be applied in human mammalian cell lines has enabled an unprecedented generation of tools^19, 20^. In this paper, we have used this technology to overcome the limitations of exogenous reporters by tagging the endogenous MYC locus with a fluorescent tag that allows for detection of MYC at the transcriptional and translational level. The MYC-EGFP signal is detected through diverse common molecular biology tools and therefore constitutes a useful tool for pooled screens and high throughput applications.

## Methods

### Cell culture and cell lines

The colorectal cancer cell lines DLD-1 and HCT-116 were maintained in DMEM (Dulbecco’s Modified Eagle Medium, Gibco) supplemented with heat-activated 10% FBS (Gibco) and 1% Penicillin/Streptomycin (Corning). Cells were maintained at 37 °C in humidified incubators with 5% CO_2_.

### CRISPR constructs

sgRNAs were chosen using the online tool Benchling. We chose three guides targeting the 20-nt area around the MYC terminal codon. For the in vitro sgRNA assay, guide RNAs were obtained from IDT. For lentiviral transduction of mammalian cells, oligos were obtained from IDT for cloning into the PX459-Puro v2.0 vector (Addgene # 62988) according to the protocol graciously shared by the F. Zhang Lab.

### HDR template constructs

pBluescript II KS (+) was digested with BamHI and NotI enzymes according to the manufacturers’ recommendations. The digest reaction was run in a 2% agarose gel and the digested plasmid was extracted using the Qiagen gel purification kit. The EGFP coding sequence was amplified from the pcDNA3-EGFP (Addgene #13031) through PCR with primers containing flanking adapters to attach BamHI and NotI sites to the fragment. Left and right homology arms of ~500bp of length were amplified through PCR. The left homology arm was amplified and BamHI restriction sites were added at both ends using appropriate primers. Similarly, the right homology arm was amplified and NotI restriction sites were added at both ends using appropriate primers. To make the construct CRISPR/Cas9-resistance, the AGG PAM target sequence was changed to AAG. The three resulting fragments were clone into the PKS digested vector in the following order 1) EGFP, 2) left arm and 3) right arm.

### Constructs for imaging

A pcDNA3.1 plasmid was digested with NotI and BsrGI according to the manufacturer’s instructions. Four different MYC-linker-EGFP fragments were cloned in to the plasmid. The different linkers were: short flexible linker (GS_8_: GCGGCCGCCggaggtagcggtggctcgagcggcggaATG), long flexible linker (GS_16_: GCGGCCGCCggaggtagcggtggctcgagcggcggaggtggatcaggtggaATG), short rigid linker (AP_8_: GCGGCCGCCGCACCTGCTCCAgcacctgctccagcaATG), long rigid linker (AP_16_: GCGGCCGCCGCACCTGCTCCAgcacctgctccagcacctgctccagcacctgctccaATG).

### Delivery methods

The PX459-Puro v2.0 vector and the HDR template were simultaneously delivered to the cells through nucleofection using the Amaxa Biosystems nucleofector II (program T030). A day after transfection cells were selected with puromycin-containing media (4 ug/mL for DLD-1 and 6 ug/mL for HCT116). After two days of selection, fresh media was added to the cells.

### PCR for fusion detection

DNA was extracted from the DLD-1 and HCT-116 post-selection cell pools through the Qiagen Blood and Tissue DNA extraction kit. PCR was performed with the following fusion-specific primers (FW-GGGATCACTCTCGGCATGG, RV-TCCTGTACTCACAAGCAGCC) and the following conditions: 95ºC for 1 min, 95ºC for 15 s, 58ºC for 10 s, 68ºC for 2 mim, for 35 cycles.

### In vitro sgRNA assay

PCR was performed around the terminal codon of the c-MYC sequence to generate a 717 bp - long fragment. The amplicon was loaded on a 2% agarose gel and purified using the QIAquick gel extraction kit. A 20 uL reaction was set up containing 100 nM of amplicon, 1uM of Cas9-sgRNA RNP complex assembled per IDT instructions. The reaction was incubated at 37ºC for 60 min and then was loaded on a 2% agarose gel.

### Western blots

A 10-cm dish of 70% confluent DLD-1 or HCT-116 cells was washed twice with cold PBS and then trypsinized with 500 uL of 0.5%Trypsin-EDTA (Gibco). Cells were spun at 1000 rpm for 3 min, resuspended and washed with cold PBS containing protease inhibitor cocktail. The washed pellet was resuspended in 50 uL 0.5% Triton-100 F-buffer followed by incubation on ice for 15 min. The samples were then centrifuged at maximum speed for 15 min at 4ºC. The resulting supernatant was isolated and protein quantification was carried out through Bradford assay per manufacturer instructions (Biorad). For SDS-PAGE, 30 ug of lysed protein from each sample was loaded on a 10% polyacrylamide gel and ran for 1h30 at 125V. Transfer to a PVDF membrane was carried out for 2h at 0.3 mA on ice. Membrane was blocked in 5% milk is TBS for 1hr. Primary incubation was carried out with antibody at 1:1000 ratio and left overnight in cold room. Secondary incubation was carried out with antibody at 1:15000 in TBS-T for 1 hr and developed in the Odyssey CRL platform.

### GFP Immunoprecipitation

Cells from an 80% confluent 10-cm plate were harvested and the immunoprecipitation protocol from Chromotek was followed. Resulting eluates and supernatants were loaded on a 10% SDS-PAGE gel and then probed with the MYC (N262) primary antibody (Santa Cruz).

### Flow cytometry

One million cells were first washed in cold PBS and then harvested in basic sorting buffer (1X PBS, 1 mM EDTA, 25 mM HEPES pH 7.0, 1%FBS). Cells were fixed with cold 70% ethanol overnight. Samples were concentrated and resuspended in sorting buffer and then analyzed for EGFP signal in a MacsQuant IML.

### Serum starvation experiment

Cells were plated in two 6-well plates and cultured in 1xDMEM (10% FBS, 1%Pen-Strep) until 80% confluent. At this point, media was changed to serum-free 1xDMEM for 48 h. Then, the first “t0” sample was harvested. Serum-free media was replaced by complete fresh media and samples were collected every hour after serum addition (t1,t2…t6). Once samples were collected and flash frozen, they were processed for western blotting.

### Cell proliferation assay

We plated 10,000 cells per well in 96-well plate. We added 10 uL of Presto Blue (Thermo Fisher) to wells read each day. After a 20 min incubation at 37ºC, absorbance measurements for 570 nm and 600 nm were taken in triplicates every day for 5 days.

### Real-time PCR

RNA was extracted from cells at same confluency via RNeasy Mini Kit (Qiagen) and DNaseI treated. cDNA was synthesized using iScript cDNA synthesis kit (Biorad). RT-PCR was set up using iQ Sybr Green Supermix (Biorad) with the appropriate primers.

## Results

### Effect of linkers on MYC-EGFP expression

EGFP reporters commonly contain the self-cleaving T2A to link both sequences together. However, we were looking to have a reporter of mRNA transcription and protein expression. Therefore, plasmid where the MYC-EGFP fusion is expressed under a constitutively active promoter were built. Each of them used different linkers to fuse the two sequences (Supplemental Fig. 1a). We transfected one of the colorectal adenocarcinoma cell lines (DLD-1) with 2 ug of plasmid and observed EGFP signal for all of the constructs and the signal was located at the nuclei. This indicated that all tested linkers allow for the expression of the MYC-EGFP fusion with some differences potentially due to uneven transfection (Supplemental Fig.1b). We also transfected the cells with a pcDNA3-EGFP and that EGFP signal was localized to the cytoplasm, as expected from the lack of an NLS sequence.

**Figure 1.**
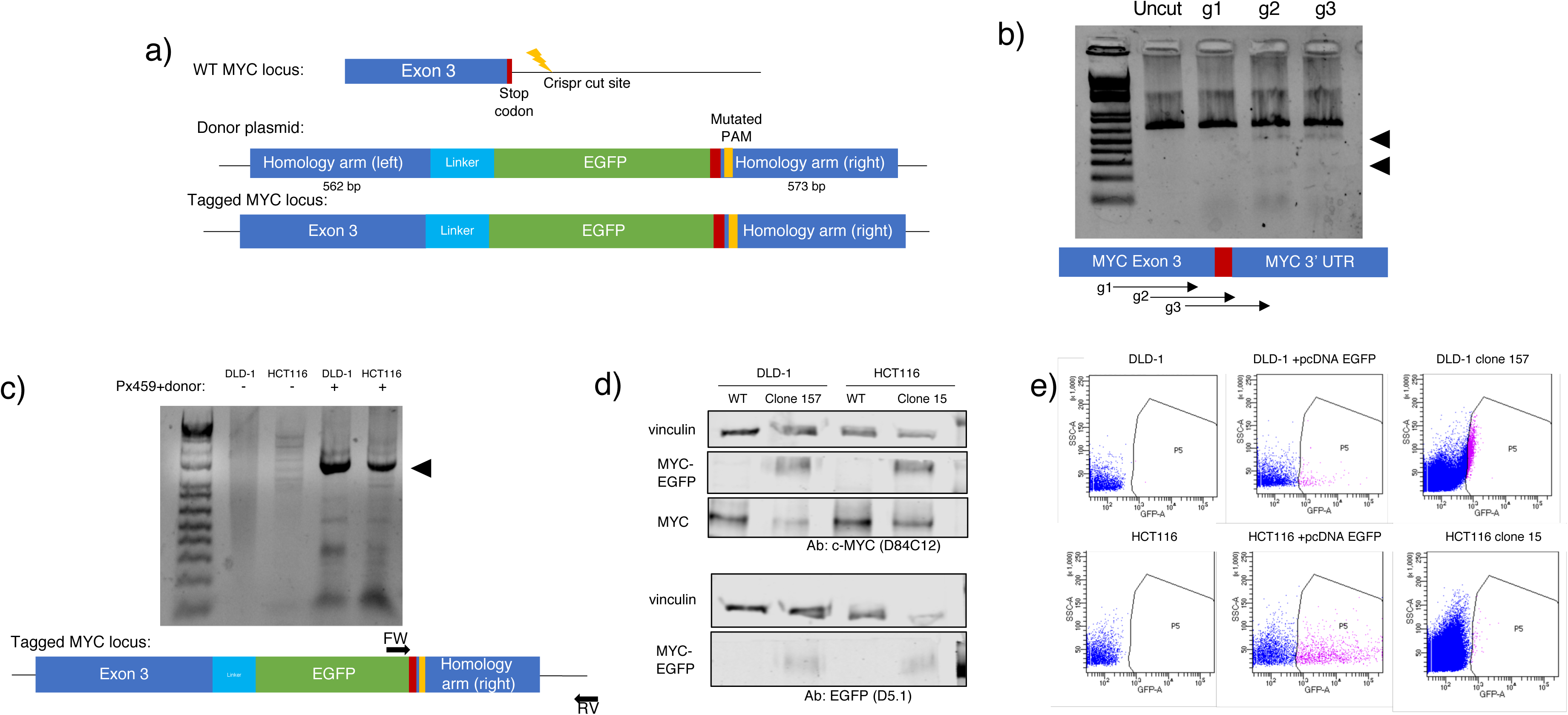
Generation of MYC-EGFP endogenous reporter in colorectal cell lines: a) General strategy for tagging exon 3 of the MYC locus through DSB mediated by CRISPR/Cas9. The donor template consists of around 500 bp homology arms, with different linkers fusing the MYC 3’ end to EGFP. A mutation changing a AGG sequence to AAG made the construct CRISPR/Cas9 resistant. b) In-vitro cleaving efficiency of three different guides was tested by digesting an amplicon with the corresponding RNPs. Cleaved products produced by each guide are seen on an agarose gel around the 200 and 400 bp marker. c) Cell lines transfected with the CRISPR lv2 vector and donor plasmid show a fusion band of 1570 bp. d) Fusion MYC-EGFP endogenous protein is readily detected through western blot, MYC WT protein is detected at 57 KDa and fusion protein at 87 KDa. e) Flow cytometry was performed on untagged cell lines, untagged cell lines transfected with pcDNA-EGFP and the tagged clonal cell lines (DLD clone 157 and HCT-116 clone 15). Endogenous EGFP signal from the tagged cell lines is effectively detected.

### Efficiency of sgRNA cleavage

CRISPR/Cas9 cleaving efficiency relies on two main aspects ^21^. The on-target efficiency is determined by the activity of a guide at the targeted region. Meanwhile, the off-target activity results from the promiscuous binding of the sgRNA to other regions, such as high homology regions followed by alternate PAM sequences. Although many algorithms provide with information about the quality of guide RNAs, we decided to carry out an in-vitro gRNA on-target cleaving assay. Three guides were selected according to their proximity to the terminal codon of the MYC protein (Fig.1b, Table 2). All guides tested were all located within 40 bp around the stop codon. A 600 bp amplicon spanning the region targeted by the guides (Table 1) was incubated with RNP complexes containing each of the three selected guides. Guide 1 was ruled out for further use due to the low level of cleavage. Guide 2 showed an evidently greater cleavage efficiency but was ruled out because CRISPR/Cas9-resistant HDR templates couldn’t be built by just changing its PAM sequence due to its effect on the end codon of MYC. Therefore, we settled for Guide 3 which showed a good amount of in-vitro cleavage (Fig 1b).

**Table 1.**
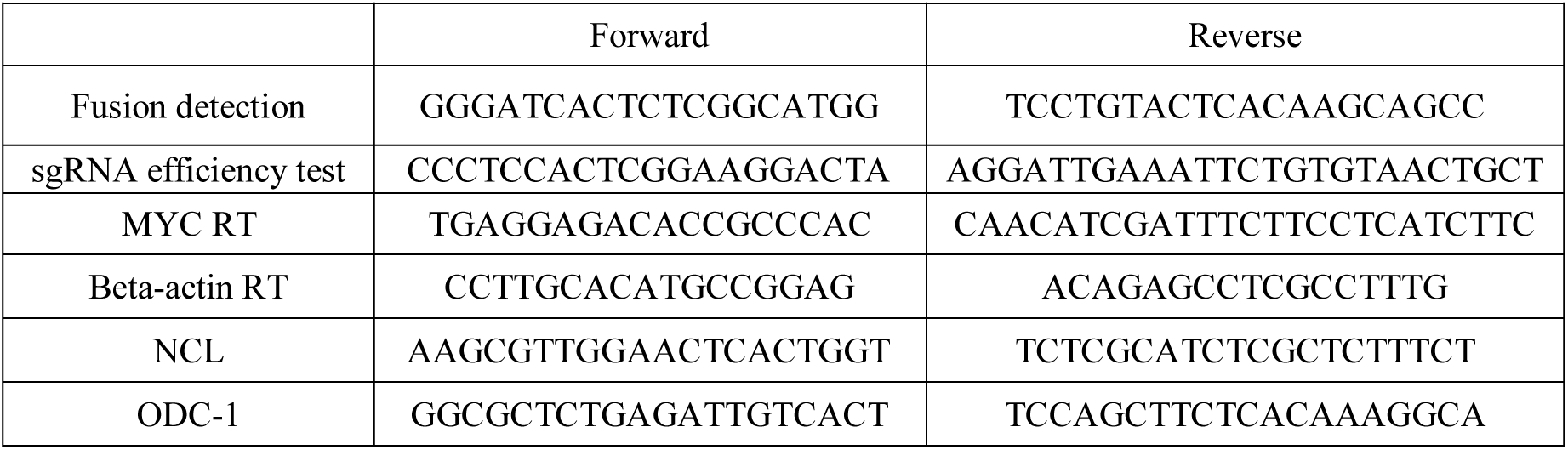
Primers used for PCR and RT-PCR.

**Table 2.**
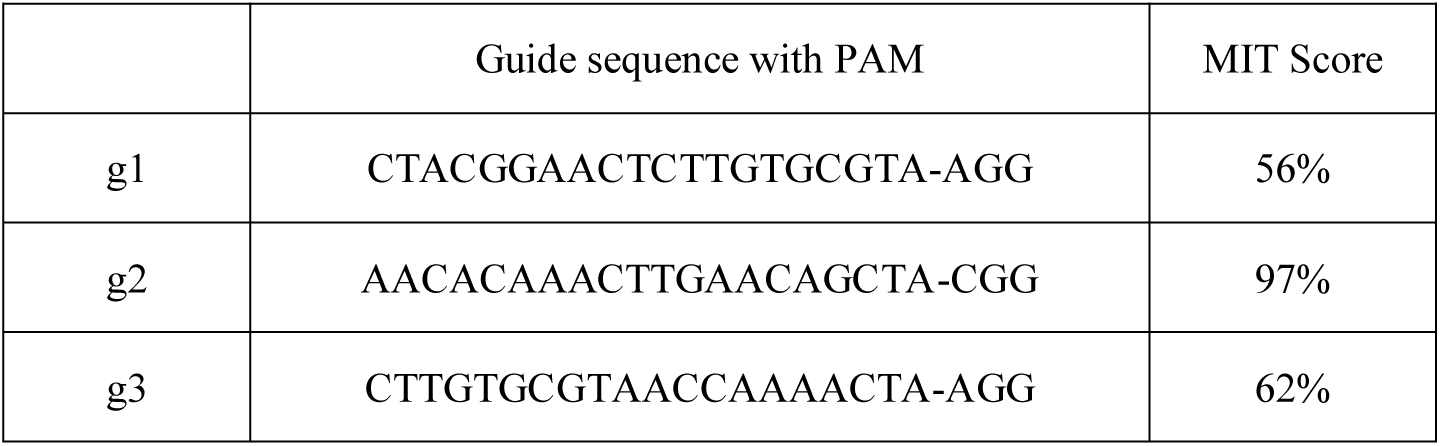
Guide sequences tested for efficiency

### PCR detection of fusion allele

In order to solely detect the presence of the fusion at the endogenous level, we designed primers that would target a sequence spanning the 3’ region of EGFP going beyond the MYC 3’ UTR region used to build one of the homology arms. Thus, these primers did not pick up the donor template. After nucleofection, PCR was carried out at 72 hr and the desired 1.5 kb product was detected in the DLD-1 and HCT-116 pools (Fig. 1c,Table 1). Then, single cells from the pools were plated in 96-well plates. Clones in each well were genotyped with the same fusion primers used for the parental pools. Positive clones were detected in both cell colorectal adenocarcinoma cell lines. In DLD-1, the HDR efficiency was 1.3% (2 out of 150 clones) and in HCT-116 the HDR efficiency was 1.25% (2 out of 160 clones). We sequenced the fusion PCR product to confirm that the expected homologous recombination had taken place and we observed that the endogenous locus showed incorporation of the CRISPR-resistant PAM sequence in DLD-1(clone 157) and HCT-116 (clone 15) (Supplementary Fig. 1c). Additionally, the area around the integrated linker was sequenced to identify which linker had been incorporated, DLD-1 (clone 157) contained a flexible linker and HCT-116 (clone 15) contained a long rigid linker (Supplementary Fig. 1c).

### MYC-EGFP fusion is detected through various techniques

Clonal populations DLD-1 (clone 157) and HCT-116 (clone 15) were expanded. MYC and MYC-EGFP fusion protein expression was confirmed through Western blot analysis of cell lysates with a polyclonal rabbit anti-MYC antibody. Additionally, a monoclonal rabbit EGFP antibody detects the fusion protein specifically (Fig. 1d). Through the use of a GFP-Trap, we confirmed that the full-length fusion expressed in the DLD-1 clone 157 and HCT116 clone 15 can be immunoprecipitated through the binding of the EGFP tag to agarose beads. The EGFP tag is not being cleaved and the full-length fusion protein runs at 90 KDa when probed with anti-MYC antibody. Meanwhile, the fusion protein is not detected in the supernatant nor eluate of the parental cell lines (Fig. 2d). We believed that the WT-EGFP fusion protein expression could function as a quick readout of MYC activity, Indeed, the EGFP signal from the endogenously tagged allele produced a fluorescent signal that was detected by flow cytometry compared to untagged parental cell line (negative control) and untagged parental cell line transfected with pcDNA3-EGFP (positive control) (Fig. 2e).

**Figure 2.**
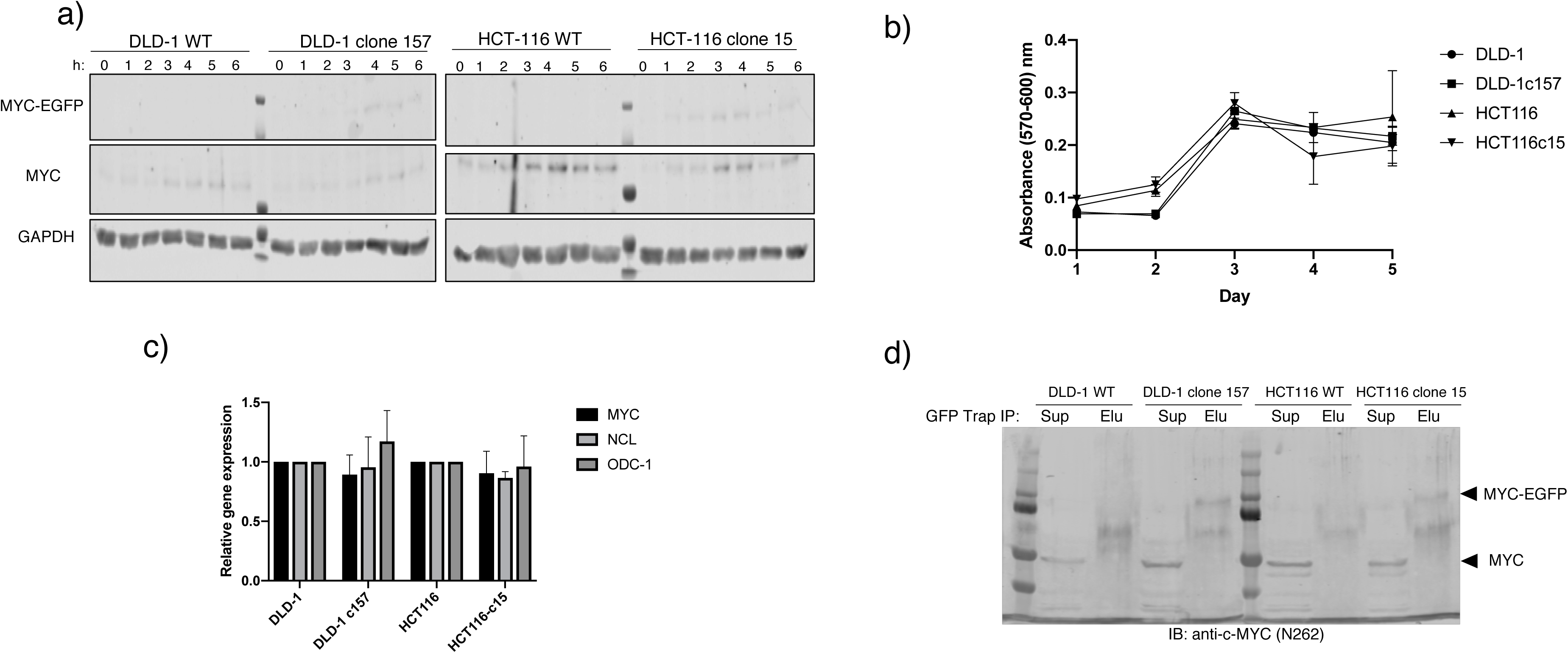
EGFP-tagged version of MYC locus is a useful readout of MYC expression: a) Serum starvation experiment: After 48-hr serum starvation, serum-containing media was added to cells and protein was harvested every hour. Blotting for MYC through western blot shows increasing production of both WT MYC (57 KDa and MYC-EGFP proteins (90 KDa) b) Cell proliferation was measured through and absorbance assay over 5 days. Parental and clonal cell lines maintain similar growth trends. c) c-MYC mRNA levels remain unchanged among parental cell lines and clonal lines. c-MYC known targets NCL and ODC-1 mRNA levels also remain the same among parental and clonal lines (levels relative to actin beta). d) Immunoprecipitation of EGFP-MYC fusion through a GFP-Trap shows pull down of fusion protein exclusively in the eluate (Elu) from tagged clones, while untagged cell lines did not. WT MYC remains in the pre-IP supernatant (Sup) of both untagged and tagged cell lines.

### MYC-EGFP fusion recapitulates the dynamics of untagged MYC in a cell

MYC expression is associated with the action of growth factors. Transfection-mediated experiments have shown that MYC exhibits serum-responsive activity. Cells were maintained in serum-starved conditions for 48 hr. Then, the media was replaced with serum-containing media and cells were harvested every hour for 6 hr after media change. Through western blot, MYC protein expression was observed and it increased consistently after addition of serum-containing media, similarly to the increase seen in parental cells lines. Additionally, we confirmed that the expression of the fusion protein follows the same dynamics as the WT MYC protein expressed in the same clone. Indeed, the proteins expressed from both MYC alleles in the tagged clones follow the same increasing trend (Fig. 2a).

### Gene expression and cell proliferation patterns are similar between tagged clonal populations and parental cell lines

Knowing that MYC expression is tightly correlated to cell cycle entry, we performed a cell proliferation curve over a period of 5 days and observed that the growth trends remain similar between the tagged clones and the corresponding parental cell line (Fig. 2b). Through RT-PCR we probed for mRNA levels of total MYC in the tagged clones and the parental cells. Total MYC levels were similar between the tagged clone and corresponding parental cell line. Similarly, we probed for mRNA levels of known MYC target genes NCL and ODC-1. The levels of both transcripts were equally abundant in the tagged clones and their corresponding parental cell lines. (Fig. 2c, Table 1).

## Discussion

In this study, we have successfully incorporated the EGFP reporter construct at the 3’ end of the MYC locus in two adenocarcinoma cell lines (DLD-1 and HCT-116) using CRISPR/Cas9 technology and monitored allelic-specific protein expression in the resulting clonal populations. Through gene editing, we can integrate a reporter in the endogenous locus of MYC which enables us to investigate endogenous gene regulation while maintaining the intact epigenetic structure and genomic architecture. MYC expression is coupled to many physiological processes, environmental cues and signals^22^. Due to this, the identification of factors orchestrating MYC regulation has been a main area of research. Early on, such research was limited to model organisms and carried out through transient transfection assays that were thought to reflect the architecture of the MYC promoter ^6^. However, the recently identified wealth of information about distal regulatory regions, the noncoding genome and the functional consequences of chromatin architecture suggests that such episomal vectors may not recapitulate the activity of the human endogenous MYC promoter. The MYC promoter is a hub of many important signals, but how they integrate and control MYC expression remains undetermined. Through a readout of transcription such as EGFP driven by the innate promoter, we believe our system provides with a real-time useful tool for exploring the regulation of the endogenous MYC locus. It has been reported than most of the regulatory control over MYC is exerted at the transcriptional level ^23^, yet there are transcriptional and post-transcriptional control mechanisms that remain identified. We believe that this system can be employed to identify such regulators. Additionally, it has been reported that in some cases, MYC dysregulation is allelic-specific in response to certain genomic changes. For example, in the KOBK 101 Burkitt lymphoma cell line, the normal WT allele remains transcriptionally silent while the MYC allele on the translocated chromosome was actively upregulated ^24^. Similarly, when an HPV region located near MYC is knocked out, MYC expression was decreased only when one particular 8q24.22 allele was affected^25^. In a similar way, the allelic-specific upregulation of MYC through chromatin looping with a distant enhancer element that contains a cancer-associated SNP (rs6983267)^26^. This SNP is present in colon cancer cells and sustains a higher preferential binding of TCF4 to the enhancer element. These studies suggest the usefulness of a system that can track the allelic-specific MYC transcription resulting from genomic alterations and distal-interactions. For decades, individual regulatory elements have been studied to reveal their effect on transcribed genes. These elements have been found to work as enhancers of transcription through their binding of transcription factors and the formation of chromatin loops that bring them in close proximity to target genes which can be nearby or distant. MYC is surrounded by a vast gene desert that contains several disease-associated SNPs that may map to potential regulator elements ^27^. The effects of these regulatory elements cannot be fully captured by the usual exogenous reporter-based experiments. The MYC downstream gene desert in a leukemia cell line to identify enhancer regions that regulate MYC expression to different levels ^28^. Although that particular study used CRISPR interference to identify the effect of these elements on MYC-dependent cell fitness, an endogenous MYC reporter offers a quick signal-based readout that can make high throughput approaches much more feasible and can offer information about intermediate phenotypes^29^. Lately, CRISPR/Cas9 pooled screens have been carried out to identify regulators of important genes through the use of reporter constructs ^30,31, 32^. These assays can assist the discovery of novel gene regulatory elements (cis-acting factors), transcription factors binding sites or exogenous regulators (transacting factors). Our results indicate that the MYC promoter-driven EGFP signal can be detected through FACS and therefore can be very useful for such high-throughput approaches targeting the coding and noncoding genome. Similarly, high throughput drug screening involves screening entire compound libraries directly against the drug target. In the era of targeted cancer therapies, gene-specific information is very valuable to avoid potential off-target-driven toxicities. Lately, drug development has focused on targeting epigenetic readers as a way of indirectly targeting cancer driver genes ^33^. A reporter system such as the one described here can be useful to discover new epigenetic targets that target MYC. The advantages of molecular reporters are innumerable and the era of CRISPR has immensely increased our ability to generate more informative biological systems.

