## Supplemental Figure 1 for "A novel tool for monitoring endogenous human MYC transcription and translation by EGFP tag insertion at the 3’ end using CRISPR-Cas9 genome editing"

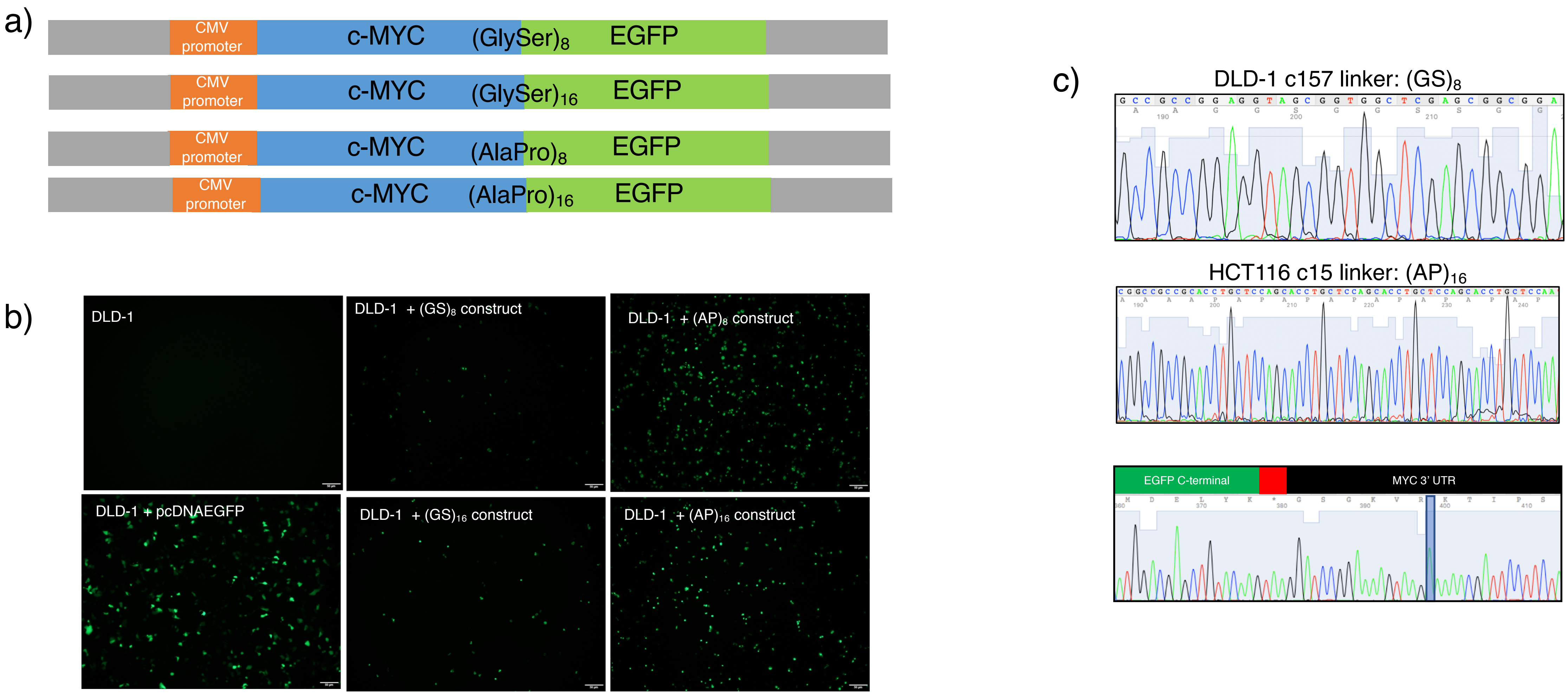


**Supplemental Figure 1. Effects of different linkers in the expression of exogenous MYC tagged with EGFP** a) To study the effect of different linkers present in the MYC-EGFP fusion, four different constructs were built on a CMV-MYC plasmid. Two flexible linkers of different length and two rigid linkers of different length were built. b) DLD-1 cells were transfected with the four different constructs. A positive control was transfected with pcDNA-EGFP and a negative control was not transfected. Images were taken at in a Olympus IX73 fluorescence microscope at 40X. c) After successful HDR in DLD-1 and HCT116 cell lines, we sequenced the region around the integrated linker. DLD-1 clone 157 shows a fusion mediated by a (GS)8 linker while HCT116 clone 15 shows a fusion mediated by a $(AP)_{16}$ linker. Sequencing of the EGFP C-terminal shows the incorporation of the tag plus CRISPR-resistant mutation (blue).
